# Isoflurane preferentially modulates synaptic responses to corticocortical stimulation over thalamocortical stimulation

**DOI:** 10.64898/2026.02.09.704944

**Authors:** Samantha Wright, Matthew I. Banks, Aeyal Raz

**Affiliations:** Department of Neuroscience, SMPH, University of Wisconsin, Madison, WI, USA; Department of Anesthesiology, SMPH, University of Wisconsin, Madison, WI, USA; Department of Anesthesiology, Rambam Healthcare Campus, Haifa, Israel; Ruth and Bruce Rappaport Faculty of Medicine, Technion – Israel Institute of Technology, Haifa, Israel

**Keywords:** Isoflurane, anesthesia, thalamocortical, cortico-cortical

## Abstract

**Objective:** To test the effect of Isoflurane on synaptic transmission of cortico-cortical and thalamocortical projections to the auditory cortex, and investigate how it modulates cortical sensory information processing to produce unconsciousness.

**Methods:** Using murine auditory thalamocortical brain slices, afferent pathways from the medial geniculate body (MGB) and layer 1 of the proximal cortex were stimulated to evoke excitatory postsynaptic potentials (eEPSPs) in cortical neurons. Whole-cell recordings were made from pyramidal and fast-spiking neurons in layer 2/3 and layer 5. eEPSPs were evaluated along with intrinsic membrane properties in response to stimulation of both pathways with and without isoflurane.

**Results:** Isoflurane administration resulted in significant eEPSP amplitude reduction following stimulation of both thalamic and cortical pathways, in layer 2/3 (p=0.015, p<0.001) and layer 5 (p<0.001, p<0.001) pyramidal neurons; while it only significantly reduced eEPSP amplitude in fast-spiking interneurons with cortical stimulation (p<0.001). Overall, isoflurane preferentially suppressed synaptic responses to cortico-cortical stimulation compared to thalamocortical (p=0.0002). Under isoflurane, cortico-cortical compared to thalamocortical stimulation evoked eEPSPs with reduced 10-90% rise time in both layer 2/3 and 5 pyramidal neurons, and shorter latency layer 5 neurons. Paired pulse ratio was not changed by isoflurane application, although an interesting loss of depression trend appear in layer 5 pyramidal neurons stimulated by cortical activation. Additional intrinsic neuronal measurements revealed that isoflurane reduced spike threshold significantly in both layer 2/3 and layer 5 neurons, reduced spike latency in layer 2/3 neurons, and input resistance in layer 5 neurons. However, these intrinsic neuronal changes were not seen in fast-spiking interneurons. All isoflurane induced changes were reversible during wash out.

**Conclusions:** Application of 1% isoflurane to brain slices significantly reduced the amplitudes of eEPSPs and modulated intrinsic neuronal properties. The effects on eEPSP amplitude were greater for cortical stimulation compared to thalamic stimulation. Isoflurane modulated intrinsic neuronal firing properties in pyramidal neurons, but not in fast-spiking interneurons.

## Introduction

How general anesthetics cause loss of consciousness is an unsolved mystery^1^. The molecular targets of anesthetic agents are widely mapped^2,3^, and the behavioral effects of these agents have been well described. However, less is known about how anesthetics act at the level of thalamocortical and cortico-cortical circuits, and in particular how anesthetics alter fundamental cortical processing features such as integration of ascending and descending information.

It has been previously shown that visual responses in A1, mediated at least in part by projections from V2^4^, are blocked by anesthetics at just-hypnotic doses that have little impact on auditory responses to pure tones^5^. These data suggest that loss of consciousness (LOC) is accompanied by reduced cortical connectivity in the presence of maintained cortical responsiveness to external stimulation. At clinically relevant doses, isoflurane significantly inhibits cortico-cortical feedback projections in both the primary auditory cortex (A1) and medial parietal association area^5,6^. At the same dose, isoflurane was found to have little impact on feed-forward and thalamocortical projections^5,6^. The preferential effect on feedback connectivity was shown to be the common feature of multiple anesthetic drugs acting on different neurotransmitter systems^7^, supporting the idea that anesthetic interference with *predictive coding* in the sensory cortex may explain LOC during anesthesia^8^.

The mechanism underlying this preferential suppression of cortico-cortical feedback pathways remains elusive. However, evaluations based on direct measures of synaptic activity are lacking, and the mechanism underlying this preferential modulation is unclear. The effects of isoflurane on the post-synaptic responses to cortico-cortical stimulation and thalamocortical stimulation recorded from primary auditory cortex pyramidal neurons are reported here. Demonstrating that the previously reported differential pathway specific effect^5^ is related to a difference in the individual post synaptic responses of pyramidal neurons. Intrinsic neuronal properties were unique between pyramidal neurons in layer 2/3 and layer 5 of the auditory cortex in response to isoflurane anesthesia.

## Methods

All experimental procedures adhered to the NIH guidelines for the care and use of Laboratory animals and were approved by the University of Wisconsin Institutional Animal Care and Use Committee.

### Slice Preparation

Thalamocortical brain slices were prepared as previously described^5,9^. Briefly, male B6CBAF1/J mice (n=58, median age=42 days old, range=28-71 days) were decapitated under isoflurane anesthesia, brains were extracted and immersed in cutting artificial CSF [cACSF: 111 mM NaCl, 35 mM NaHCO_3_, 20 mM HEPES, 1.8 mM KCl, 1.05 mM CaCl_2_, 2.8 mM MgSO_4_, 1.2 mM KH_2_PO_4_, and 10 mM glucose] at 0–4^°^C. Auditory Thalamic-Cortical (TC) brain slices (500 μm) were prepared from the right hemisphere^10,11^. Slices were maintained in cACSF saturated with 95%O2/5% CO2 at 24^°^C for >30 minutes before transfer to the recording chamber, which was perfused at 5–6 ml/min with ACSF [111 mM NaCl, 35 mM NaHCO3, 20 mM HEPES, 1.8 mM KCl, 2.1 mM CaCl2, 1.4 mM MgSO4, 1.2 mM KH2PO4, and 10 mM glucose] at 32– 34^°^C, heated by an inline heater (Warner Instruments model SH-27B and controller TC-324B) or via conducting gel surrounding the inlet perfusion system (controlled by LN Temperatcontoler III).

### Electrophysiological Recordings

Auditory cortex was identified as previously described^12^ (Krause et al. 2014, Verbny et al., 2006; Banks et al., 2011). To electrically activate the Auditory cortex, thalamic afferents were stimulated using a pair of tungsten electrodes (0.1MW, 75μm diameter; FHC Inc., Bowdoin, ME) cemented together at tip separations of ∼50– 200μm. Stimuli (100μs, 30–250μA) were applied using constant current stimulus isolation units STG 4002, Multichannel Systems, Reutlingen, Germany) and consisted of brief trains (4 pulses at 40Hz). For activation of the auditory thalamic fibers in brain slices, stimulation was given rostral to the MGBv in the fiber bundle called the superior thalamic radiation, which runs from the auditory thalamus to the auditory cortex. In all experiments, verification of a L4 current sink in response to stimulation of the fiber bundle was made. However, the possibility that some of the recordings were outside primary auditory cortex exists. These recording would include responses from auditory areas outside of A1 that receive driving core auditory thalamic inputs, e.g., AuV or AuD in the Paxinos (Paxinos and Franklin, 2003) terminology. If at first an extracellular response was not elicited, the recording pipette was moved approximately 200-300mm rostral or caudal until a response was obtained. Extra cellular field potentials were recorded with borosilicate glass (similar composition to the whole-cell recording pipettes which are described in the following paragraph), the tips of the pipettes were broken under visual control to an outer tip diameter of 10–15μm and had open-tip resistances of about 0.5-0.75MΩ when filled with ACSF.

Whole-cell current clamp recordings were then made in the region of the extracellular responses of L4 current sinks, in either L2/3 or L5. Cortical layers were identified by differences in cell density and based on distance from the pia in conjunction with previous studies (Banks et al., 2011). Tissue appearance under bright field illumination was further used to identify the approximate borders between cortical layers. Layers 4 had a relatively dark appearance compared to the lighter colored bands of layers 3, 5, and 6. Electrophysiological recordings were obtained using patch pipettes fabricated from borosilicate glass (KG-33; 1.7mm outer diameter; 1.1mm inner diameter; Garner Glass, Claremont, CA) and pulled to final tip resistances with a Flaming-Brown two-stage puller (P-87; Sutter Instruments, Novato, CA). For whole-cell current clamp recordings, pipettes had open tip resistances of 4–7MΩ when filled with intra cellular solution (in mM):140 K-gluconate, 10 NaCl, 10 HEPES, 0.1 EGTA, 4 Mg ATP and 0.3 Na GTP, pH7.2. In some experiments 0.3% biocytin was included in the intracellular solution for post hoc anatomical identification of recorded neurons. Cells were visualized using a video camera (Hamamatsu or Q Imaging Rolera) connected to an upright microscope (Olympus BX51-WI) with a long working-distance water-immersion objective (Olympus 40X, 0.9 N.A.) and differential interference contrast optics.

While recording in whole-cell configuration current clamp, intracellular recordings were made in response to a series of hyperpolarizing and depolarizing current steps that were recorded in addition to both evoked thalamic stimulation and cortical stimulation by a second pair of tungsten electrodes stimulating extracellularly in L1 approximately 500-1000mm anterior to the recording pipette in either L2/3 or L5. After a control set of recordings were made, Isoflurane dissolved in ACSF was perfused over the slice for 5-10 minutes and the same series of recordings was made, followed by a wash out period of approximately 10-20 minutes of control ACSF where the final series of recordings were made.

### Isoflurane Preparation

Isoflurane (Piramal Healthcare) was dissolved in ACSF as an aqueous solution in a Teflon gas sampling bag (Fisher Scientific International Inc). Approximately 30ml of isoflurane air gas from this bag was mixed with approximately 270ml of saturated 95%O2-5%CO2 in a second Teflon gas sampling bag. The concentration of isoflurane was measured to be at approximately 3.3%; where from approximately 180ml of this stock gas solution was mixed in a third Teflon gas sampling bag with approximately 120ml of saturated 95%O2-5%CO2 and mixed with 300ml ACSF for a 50%-50% gas/ACSF final volume. The control bag of ACSF was 300ml saturated 95%O2-5%CO2 mixed with 300ml ACSF. Both bags were mixed on a shaker table for at least 1 hour but not longer than 24 hours. The final concentration of isoflurane was verified by measuring gas concentration at equilibrium with gas in the bag after experiments using an anesthetic gas monitor (Poet II Anesthesia Monitor Criticare Systems Inc.) and was set around 1% of isoflurane for experimental perfusion.

### Data Acquisition

Data were amplified (MultiClamp-700A; Molecular Devices, Union City, CA), low-pass filtered (10 kHz), digitized at 20 kHz and 40kHz (Digidata 1440A; Molecular Devices, Sunnyvale, CA) and recorded using pClamp version 9.2 and 10.6 (Molecular Devices).

### Data Analysis

Excitatory postsynaptic potential (EPSP) amplitude, 10-90% rise time, latency and ratio were measured from the average of ten responses to the first response from a train of 4 pulses delivered at 40 Hz. EPSP amplitude was measured as rest-to-peak and EPSP rise time was measured as the time from 10% to 90% of peak on the rising phase of the evoked potential. EPSP latencies were measured as the time of rise to 10% of the peak. EPSP ratio was the amplitude of the first evoked EPSP from the isoflurane and wash conditions normalized to the control amplitude and are reported in percentage of the magnitude of the first response of the control. In whole-cell recordings, contribution from inhibitory postsynaptic potential (IPSP) during evoked stimulation from the thalamus and cortex was identified by holding the cells at depolarized potentials to spike threshold. If an inhibitory component was observed in the evoked EPSP, the recording was excluded from EPSP analyses.

Intrinsic cellular properties were measured from eleven sweeps of hyperpolarizing and depolarizing current steps in whole-cell configuration. Resting membrane potential was averaged from the sweeps before current injection, input resistance was measured by V=I/R at the smallest depolarizing and hyperpolarizing current steps. Spike latency was calculated from the time current was injected until time of first spike (measured as the 2^nd^ derivative) in the lowest current step which elicited a spike. Spike threshold was based on the 2^nd^ derivative of the first spike elicited by the lowest amplitude current step which evoked an action potential. The time constant was calculated as the time it took the membrane potential to return to 2/3^rds^ of the resting state value after current was injected.

### Statistical Methods

Statistical analysis for the intrinsic cellular properties and synaptic currents, compared between control, isoflurane and wash saline, were performed in SPSS using a Generalized Linear Model with repeated measures analysis of variance. A p-value of less than 0.05 is regarded as significant. For each metric that was assed statistically, a custom code was written in MATLAB to extract the numerical values. In the measurements of postsynaptic currents, multiple sweeps during a single recording were averaged for the analysis.

To compare the statistical significance between the reduction of the eEPSPs by isoflurane from thalamic stimulation versus cortical stimulation a Linear Mixed – Effects Model fit by ML was used with a 95% confidence interval. Using a theoretical likelihood ratio test comparing:

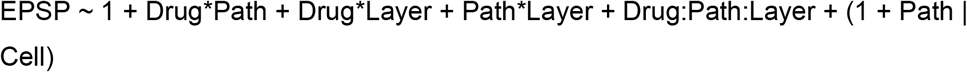

To

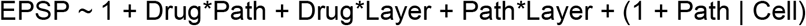

As well as a reduced model of Theoretical Likelihood Ratio Test comparing:

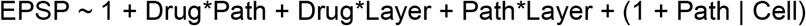

To

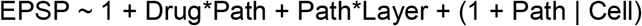

This formula allowed determination of the significant effect of isoflurane on thalamocortical versus corticocortical eEPSPs.

## Results

Brain slice preparations were cut with an orientation that preserves the fiber bundle from the medial geniculate body (MGV) to the auditory cortex (AC), and whole-cell patch-clamp recordings from neurons in layer 2/3 and layer 5 of the AC were made. Stimulation of the thalamocortical (TC) pathway via the fiber bundle from the MGV (noted with the double asterisks, on top in **Figure 1A**), as well as cortico-cortical (CC) pathway at layer 1 of the proximal cortex (double asterisk on the bottom of **Figure 1A**), was made using a train of 4 stimuli at 40 Hz. In **Figure 1A**, the recording pipette is illustrated with a single asterisk on the bottom left. **Figure 1B** illustrates an eEPSP, and demonstrates the analyzed parameters: 10-90% rise time (green circles) and peak of the eEPSP (red circle) were measured in three conditions: control (dark gray trace), isoflurane (dashed trace) and wash (light gray trace). Additionally, latency to the onset of the eEPSP was assessed. Each metric was quantified following stimulation of both pathways, in all conditions: control, isoflurane, and wash. The mean values were calculated from the average of 11 eEPSP sweeps.

**Figure 1.**
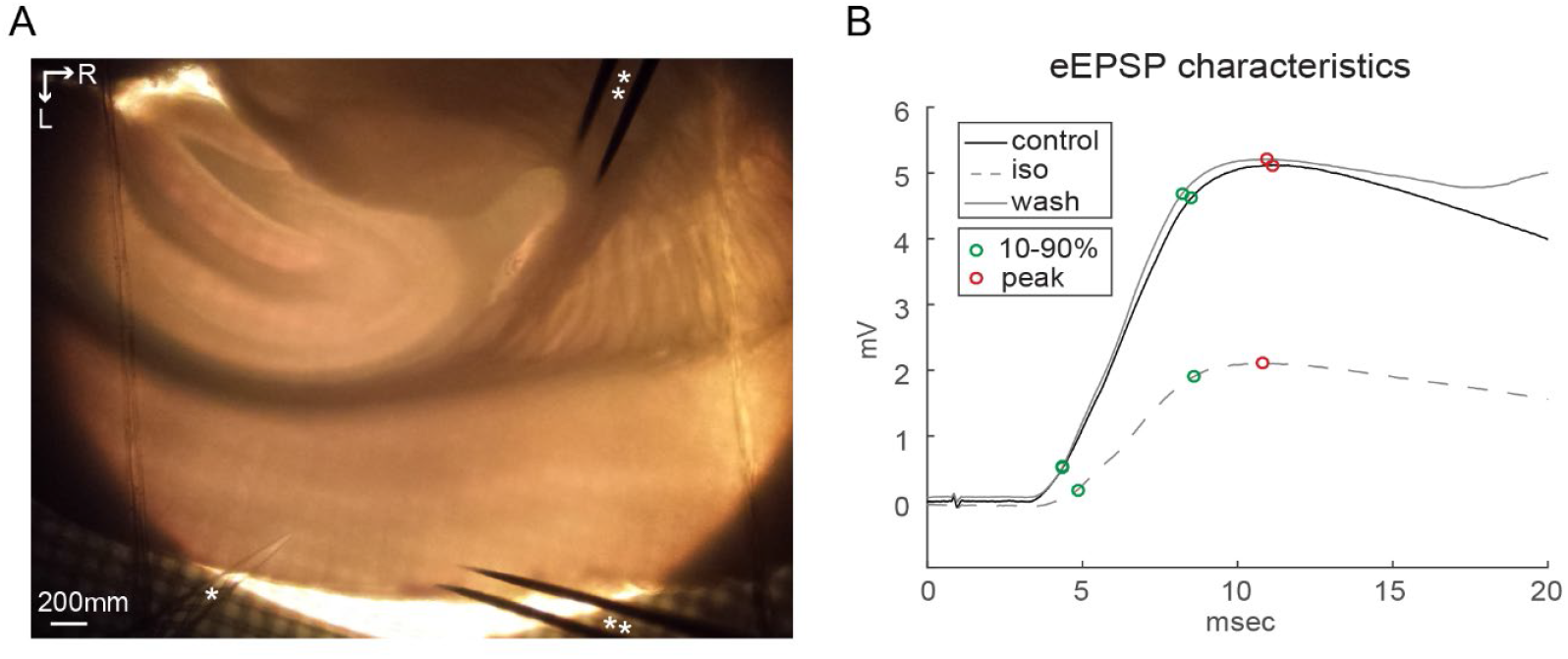
Thalamocortical Brain Slice and eEPSP Characteristics. (A)500 μm thick murine brain slice containing the auditory cortex (AC), the ventral division of the medial geniculate body (MGv) and the fiber bundle connecting them. Single asterisk denotes the recording pipette and double asterisks denotes the stimulating electrodes in the AC and thalamic fiber bundle. (B)Representative eEPSPs recorded from a pyramidal neuron in the AC in response to electrical stimulation under control saline (black), isoflurane saline (gray dash) and wash saline (gray) conditions. Peak eEPSP is marked by the red circles, 10% to 90% rise of the eEPSP are marked between the green circles. Latency measurements were made from the time between stimulus and positive deflection of the eEPSP (not shown).

Pyramidal and fast-spiking neurons in layers 2/3 and 5 were distinguished by their differential spiking responses to hyperpolarizing current injection, as well as shape of the soma. Representative whole-cell recordings are shown in **Figure 2 A-C** from a pyramidal neuron in layer 2/3 in control, isoflurane and wash conditions, respectively. Similarly, **Figure 2 D-F** shows a representative layer 5 pyramidal neuron recorded with all conditions, as well as a fast-spiking interneuron in **figure 2 G-I**. To quantify how isoflurane affected neuronal spiking, several intrinsic cellular firing characteristics including resting membrane potential (**figure 3A**), spike latency (**figure 3B**), spike threshold (**figure 3C**), input resistance (**figure 3D**), and membrane time constant (**figure 3E**) were analyzed. Examples showing where spike threshold was measured are illustrated in a layer 2/3 pyramidal neuron (**figure 3F**), and a layer 5 pyramidal neuron (**figure 3G**).

**Figure 2.**
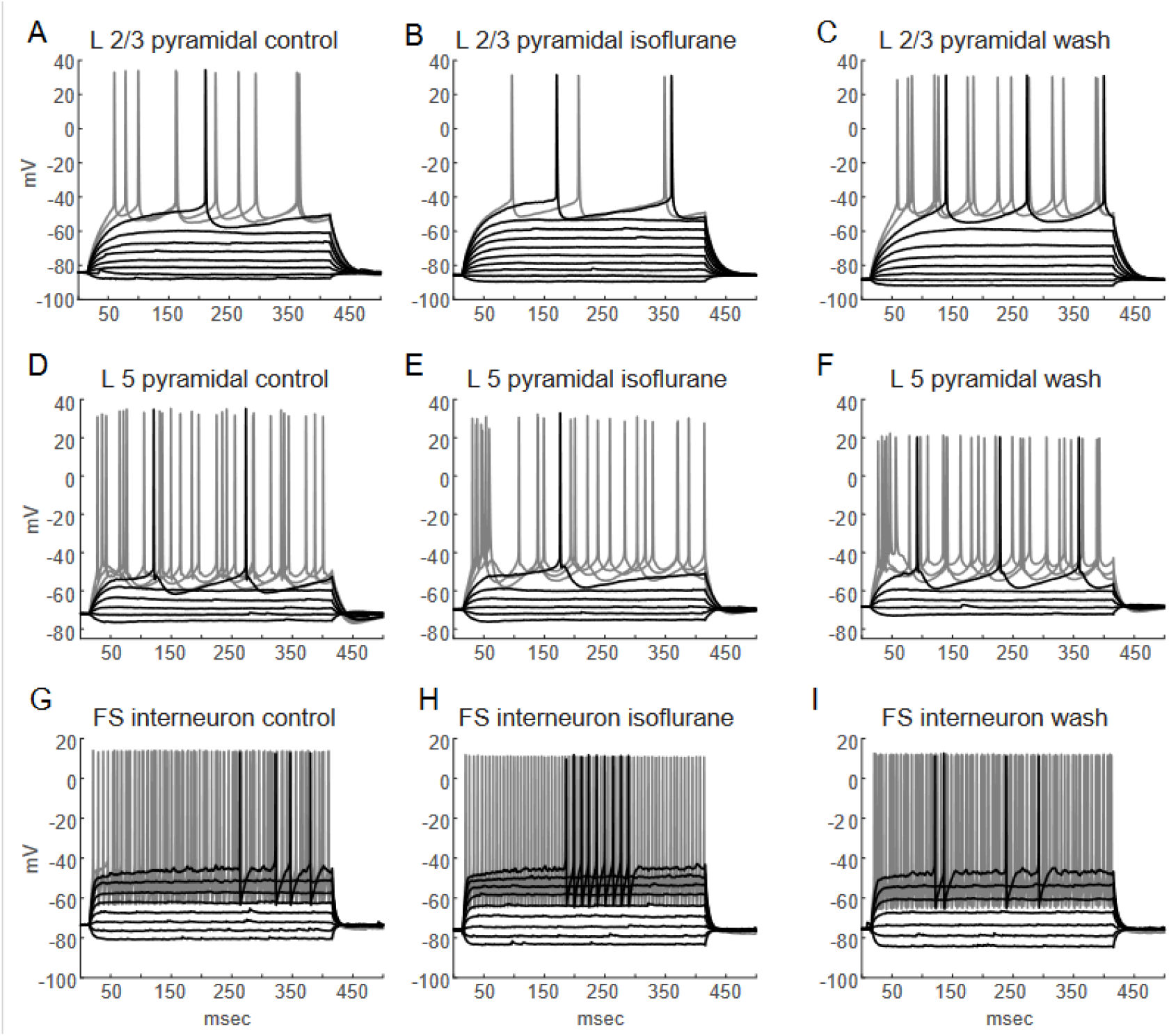
Electrical Signatures of Cortical Neurons in Response to Isoflurane. (A-C) Layer 2/3 pyramidal neuron recorded in whole-cell configuration in response to current steps with control saline (A), isoflurane saline (B) and wash saline (C). (D-F) Layer 5 pyramidal neuron recorded in whole-cell configuration in response to current steps with control saline (D), isoflurane saline (E) and wash saline (F). (G-I) Fast Spiking interneuron recorded in whole-cell configuration in response to current steps with control saline (G), isoflurane saline (H) and wash saline (I).

**Figure 3.**
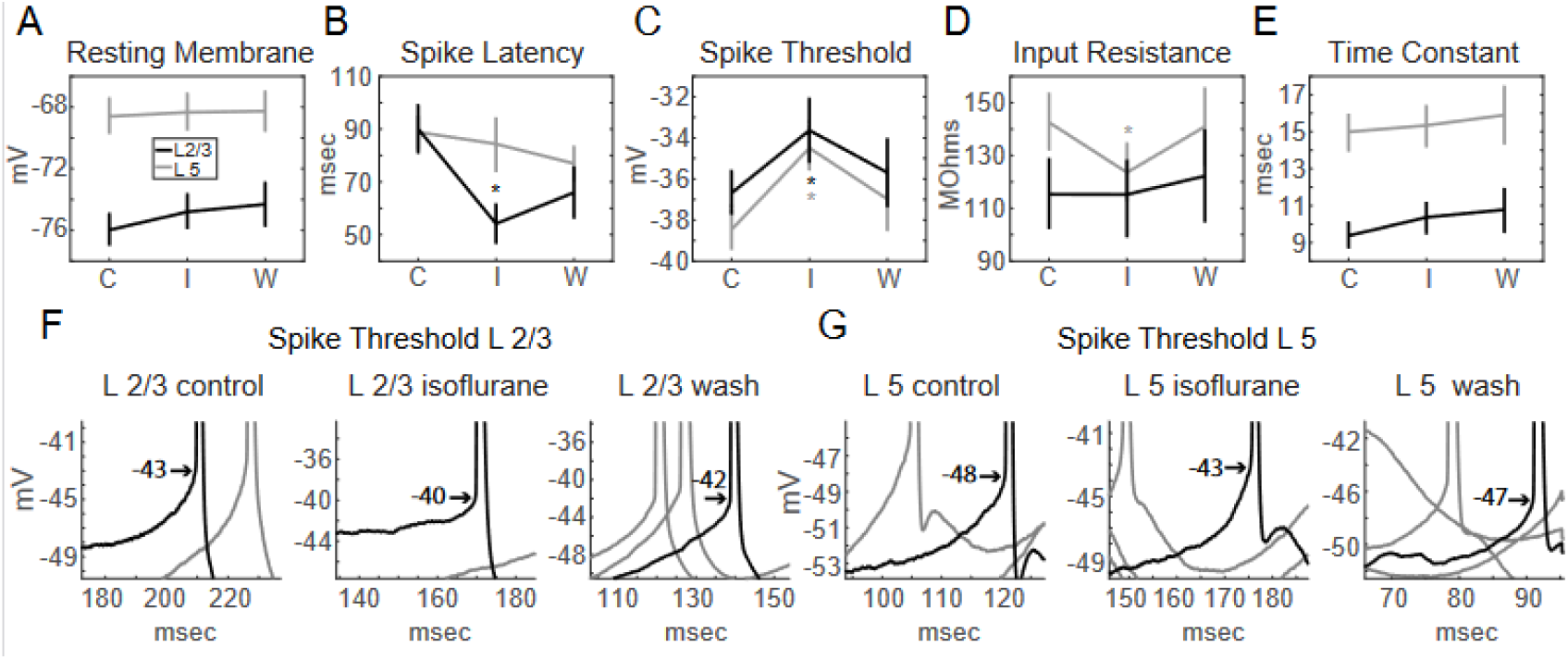
Intrinsic Cellular Properties of Pyramidal Neurons Affected by Isoflurane. (A)Averaged resting membrane potential of layer 2/3 (black) and layer 5 (gray) pyramidal neurons was not significantly affected by isoflurane saline (I) as compared to control (C) and wash (W) saline. (B)Averaged spike latency, time from current injection to the first spike at the lowest current step which elicited and action potential, was significantly faster in layer 5 neurons (gray trace) under isoflurane conditions (I) as compared to control saline (C) and wash conditions (W). In layer 2/3 neurons (black trace) the spike latency was not significantly affected by isoflurane. (C)Averaged spike threshold was significantly depolarized by isoflurane (I) in both layer 2/3 (black) and layer 5 (gray) pyramidal neurons, as compared to control saline (C)and wash saline (W). (D)Averaged input resistance was significantly reduced by isoflurane (I) as compared to control saline (C) and wash conditions (W) in layer 5 pyramidal neurons (gray), but not in layer 2/3 neurons (black trace). (E)Averaged membrane time constant of pyramidal neurons was not significantly affected by isoflurane application (I) compared to control saline (C) and wash saline (W) in either layer 2/3 nor layer 5 pyramidal neurons. (F)Representative layer 2/3 pyramidal neuron, with a more depolarized spike threshold under isoflurane saline conditions (middle) compared to control and wash saline (left and right, respectively), arrows illustrate where spike threshold was measured as the second derivative. (G)Representative layer 5 pyramidal neuron, with a more depolarized spike threshold under isoflurane saline conditions (middle) compared to control and wash saline (left and right, respectively), arrows illustrate where spike threshold was measured as the second derivative.

Each cell type was assessed individually using a generalized linear model. Some cellular properties changed significantly with isoflurane. Interestingly no significant changes during isoflurane application were seen in the intrinsic properties of fast spiking interneurons, although the number of these recordings were few as pyramidal neurons were targeted for recording. Both resting membrane potential (**3A**) and membrane time constant (**3E**) were unaffected by isoflurane in all cell types. In layer 2/3 pyramidal neurons only, spike latency (**3B**) was significantly decreased with isoflurane application (control =90.2 ms, Iso=55.0 ms, P=0.004, n=25; spike latency was approximately shorter 60% with isoflurane as to control, on average). Additionally, in layer 2/3 pyramidal neurons, spike threshold (**3C**) was significantly depolarized under isoflurane compared to control saline (control =-36.85mV, Iso=-33.62mV, P=0.003, n=26; firing around 9% more depolarized under isoflurane, on average). In layer 5 pyramidal neurons spike threshold was also significantly depolarized under isoflurane conditions (control =-38.57mV, Iso=-34.80mV, P<0.001, n=33; around10% more depolarized under isoflurane, on average). Layer 5 pyramidal neurons also had their input resistance significantly decreased by isoflurane as compared to control (control =142.82MOhms, Iso=123.52MOhms, P=0.013, n=33; reduced to approximately 86% of control under isoflurane, on average) (**3D**).

EPSPs were evoked in response to stimulus trains, induced by both thalamocortical and cortico-cortical stimulation while recording in layer 2/3 and layer 5 neurons in the AC. Representative traces are shown in **figure 4**. The stimulation paradigm included 11 sweeps of 4 pulses at 40 Hz, which was used to assess paired pulse ratio and trends of synaptic depression as well as facilitation. The sweeps were averaged for analysis. The amplitude of eEPSPs was taken from the 1^st^ response as measured from baseline to the peak of the eEPSP. **Figure 4 A-C** illustrates the individual (gray) and averaged (black) neuronal responses from a layer 2/3 pyramidal neuron via cortical stimulation from layer 1 of the cortex, adjacent to the AC. **Figure 4 D-F** is an overlay of the averaged eEPSP responses following cortical stimulation in a layer 2/3 neuron (**figure 4D**), layer 5 pyramidal neuron (**figure 4E**), and a fast-spiking interneuron (**figure 4F**) in control (black trace), isoflurane (dashed gray trace), and wash (solid gray trace) conditions. Similarly **figure 4 G-H**, illustrates responses to thalamic stimulation in the same neurons: layer 2/3 neuron (**4G**), layer 5 pyramidal neuron (**4H**), and a fast-spiking interneuron (**4I**) in all conditions.

**Figure 4.**
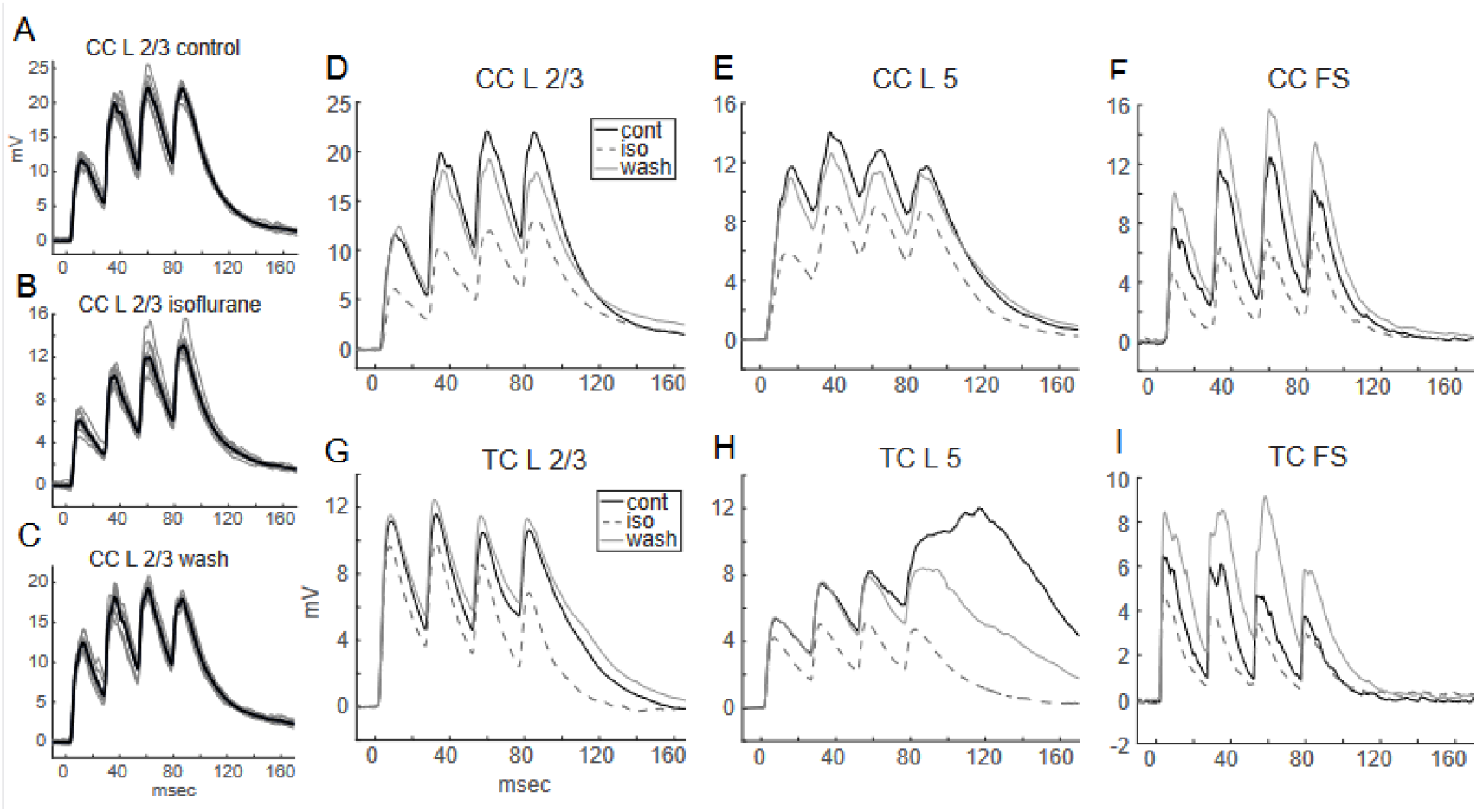
Stimulus Train of eEPSP at 40 Hz. (A-C) Layer 1 cortico-cortical stimulation inducing eEPSPs in a layer 2/3 pyramidal neuron, measured under control (A), isoflurane (B), and wash (C). Individual sweeps in gray, and averaged in black. (D)Averaged eEPSPs in a layer 2/3 pyramidal neuron evoked by cortical stimulation, in control (black), isoflurane (gray dash), and wash (gray) saline conditions. (E)Averaged eEPSPs in a layer 5 pyramidal neuron evoked by cortical stimulation, in control (black), isoflurane (gray dash), and wash (gray) saline conditions. (F)Averaged eEPSPs in a fast-spiking interneuron evoked by cortical stimulation, in control (black), isoflurane (gray dash), and wash (gray) saline conditions. (G)Averaged eEPSPs in a layer 2/3 pyramidal neuron evoked by thalamic stimulation, in control (black), isoflurane (gray dash), and wash (gray) saline conditions. (H)Averaged eEPSPs in a layer 5 pyramidal neuron evoked by thalamic stimulation, in control (black), isoflurane (gray dash), and wash (gray) saline conditions. (I)Averaged eEPSPs in a fast-spiking interneuron evoked by thalamic stimulation, in control (black), isoflurane (gray dash), and wash (gray) saline conditions.

Changes in eEPSP response during isoflurane application following cortical and thalamic stimulation in pyramidal neurons in layers 2/3 and 5 of the AC were identified. Different magnitudes of eEPSP suppression were found under isoflurane within each layer and pathway. **Figure 5** illustrates the dynamics of individual neuronal responses as well as the average, per layer and pathway. Using the averaged first eEPSP response to the stimuli, response dynamics were compared during isoflurane administration in all layers and pathways. There was a significant reduction of eEPSP amplitude by isoflurane in all pathways and layers. The individual and averaged eEPSP response to thalamic stimulation for layer 2/3 and 5 are respectively shown in **figures 5A & B**; while the individual and averaged eEPSP responses to cortical stimulation for layer 2/3 and 5 are respectively shown in **figures 5C & D**.

**Figure 5.**
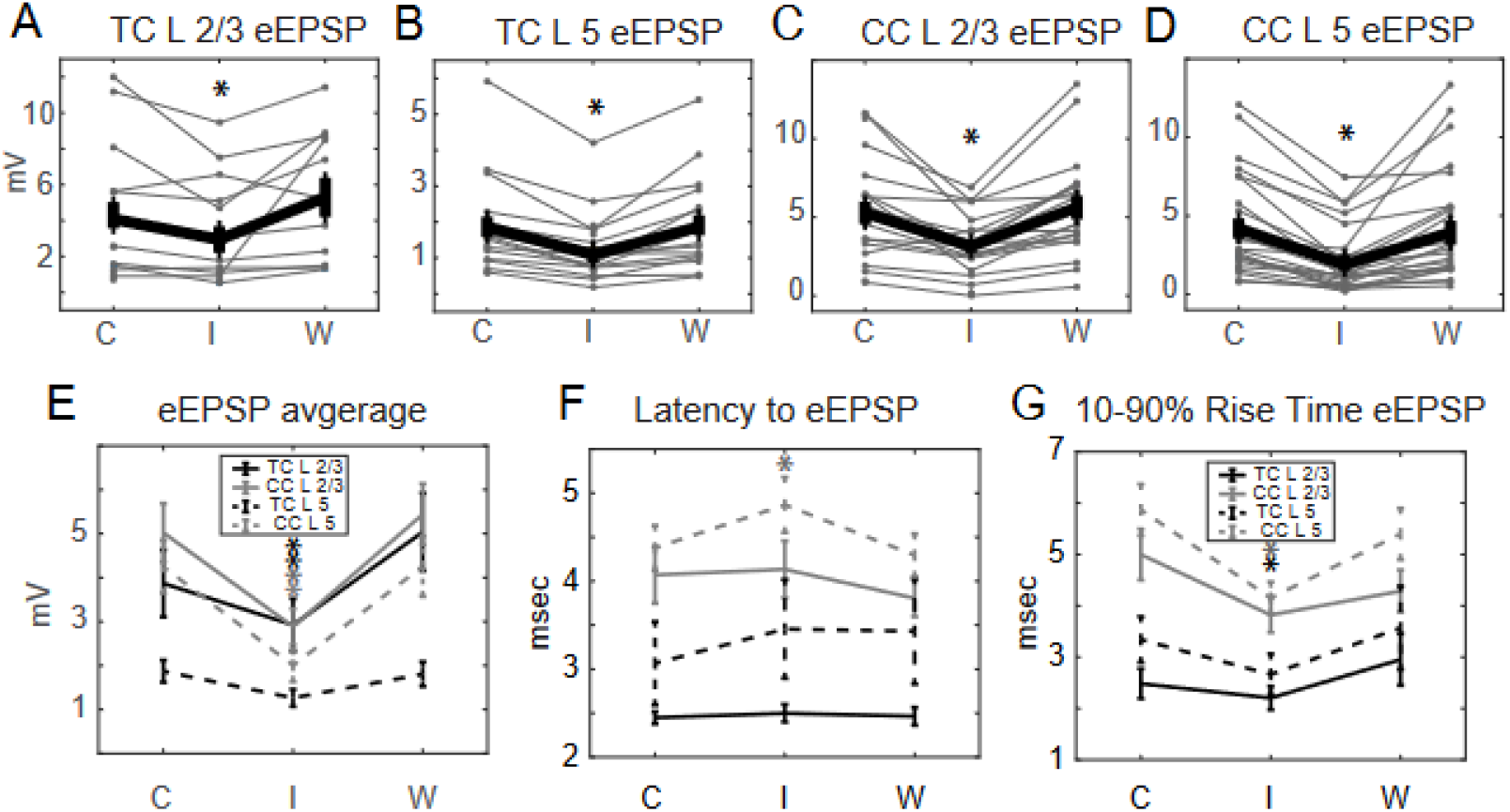
Comparison of eEPSPs Reduced by Isoflurane. (A-D) Individual (gray) and averaged (black) measurements of eEPSP amplitude recorded in pyramidal neurons in control (C), isoflurane (I) and wash (W) saline. Isoflurane significantly reduced the average eEPSP amplitude compared to that in control saline in all pathways and layers. (A) Thalamic stimulation of layer 2/3 pyramidal neurons, (B) Thalamic stimulation of layer 5 pyramidal neurons, (C) Cortical stimulation of layer 2/3 pyramidal neurons, (D) Cortical stimulation of layer 5 pyramidal neurons. (E)Mean eEPSP amplitude, significantly reduced by isoflurane in layer 2/3 and layer 5 pyramidal neurons from both from thalamic and cortical stimulation, in control (C), isoflurane (I) and wash (W) saline. Black solid line is thalamic stimulation to L 2/3, and the solid grey line is cortical stimulation to L 2/3; black dashed line is cortical stimulation to L 5, and dashed grey line is cortical stimulation to L 5. (F)Average latency from stimulation time to onset of first eEPSP. Isoflurane (I) altered eEPSP latency as compared to control (C) and wash (W) saline conditions. Cortical stimulation to layer 5 pyramidal neurons was significantly increased/delayed by isoflurane. Both cortical stimulation to layer 2/3 neurons and thalamic stimulation to layer 5 neurons neared significance with isoflurane. Black solid line shows thalamic stimulation to L 2/3, while solid grey line is cortical stimulation to L 2/3; black dashed line shows cortical stimulation to L 5, and dashed grey line is cortical stimulation to L 5. (G)Average 10-90% rise time of 1^st^ eEPSP. Isoflurane (I) application affected rise time compared to control (C) and wash (W) saline conditions. Both cortical and thalamic stimulation to layer 5 neurons (but not layer 2/3 neurons) significantly decreased rise time. Black solid line shows thalamic stimulation to L 2/3, and solid grey line shows cortical stimulation to L 2/3; black dashed line shows cortical stimulation to L 5, with dashed grey line shows cortical stimulation to L 5.

Layer 2/3 pyramidal neuron eEPSP amplitudes were significantly reduced during isoflurane application, which recovered during washout. Following thalamic stimulation EPSP amplitude under isoflurane was reduced to 77% of control (control =4.40mV, Iso=3.37mV, P=0.015, n=15) and following cortical stimulation the amplitude reduced to 60% (control = 5.32mV, Iso=3.21mV, P<0.001, n=21). The eEPSP amplitudes of Layer 5 pyramidal neurons were also significantly reduced by isoflurane, and recovered during washout. Isoflurane reduced the responses to thalamic stimulation to 64% of control (control =1.82mV, Iso=1.16mV, P<0.001, n=20) and cortical stimulation to 47% of control (control =4.42mV, Iso=2.08mV, P<0.001, n=26). **Figure 5E** shows the overlay of averaged eEPSP amplitudes from pyramidal neuron in both layers, stimulated by both pathways, as analyzed by a generalized linear model.

In layer 2/3 pyramidal neurons, significant changes during isoflurane administration were found in the eEPSP amplitude from both stimulation pathways, as well as rise time from cortical stimulation. In layer 2/3 neurons, eEPSP rise time following cortical stimulation was significantly decreased to 76% of control by isoflurane application (control=5.0ms, Iso=3.8ms, P=0.021, n=20). Response latency, as well as rise time following thalamic stimulation in layer 2/3 pyramidal neurons was not significantly changed during isoflurane application.

In layer 5 pyramidal neurons, isoflurane application altered average eEPSP rise time from TC stimulation reducing it to 80% of control under isoflurane, which approached statistical significance (control=3.5ms, Iso=2.8ms, P=0.050, n=15). Rise time of eEPSPs following CC stimulation was significantly reduced to 70% of control (control=5.7ms, Iso=4.0ms, P<0.001, n=23). Layer 5 eEPSP latency was significantly increased to 112% of control following CC stimulation (control=4.2ms, Iso=4.7ms, P=0.029, n=23, while latency following TC stimulation was not significantly altered by isoflurane. The averaged traces of latency to the first eEPSP, and 10-90% rise time are shown overlaid in **figures 5F & 5G**.

In fast spiking interneurons eEPSPs evoked by CC stimulation were significantly reduced to 55% of control by isoflurane application, recovering during washout (control=6.88mV, Iso=3.81mV, P<0.001, n=8). No other metrics in fast spiking interneurons were significantly altered by isoflurane anesthesia.

All parameters were measured separately for each of the 3 neuronal types dived into groups established by stimulation pathway independently. Additional analysis was performed by combining pathways for each layer, and by combining both cortical layers from each pathway. This structure of analyses allowed for comparison of both pathways’ effects on each layer, and for all neurons in a layer stimulated by each individual pathway. This setup revealed which aspects of the connectivity were affected by isoflurane. Many eEPSP features following TC stimulation in layers 2/3 and 5 neurons were significantly affected by isoflurane; although these alterations were often not as robust and intense as those following CC stimulation. Pyramidal neurons in both layers following stimulation via the TC pathway, had significantly reduced eEPSP amplitudes during isoflurane application (control=2.92mV, Iso=2.11mV, P=0.001, n=35). On average, eEPSP amplitudes under isoflurane were 72% of control. Both rise time and latency were statistically unaffected by isoflurane with thalamic stimulation.

Stimulation of the CC pathway was significantly altered by isoflurane, often with a greater effect than the responses to thalamic stimulation. Pyramidal neurons in both layers demonstrated significantly reduced amplitudes during isoflurane application (control=4.82mV, Iso=2.58mV, P<0.001, n=47). The average eEPSP amplitude following CC stimulation was reduced to 54% of control under isoflurane. Isoflurane affected CC triggered eEPSP rise time (control=5.3ms, Iso=3.9ms, P<0.001, n=43), which was reduced to 74% of control under isoflurane, and latency (control=4.0ms, Iso=4.3ms, P=0.028, n=43), which increased to 108% of control under isoflurane.

Similarly, combining all responses recorded in layer 2/3 (following stimulation of both pathways) demonstrated significant reductions in most eEPSP features during isoflurane application, except latency, which was unchanged by isoflurane. Stimulation of pyramidal neurons in layer 2/3 led to a significantly reduced eEPSP amplitudes during isoflurane application (control=4.94mV, Iso=3.28mV, P<0.001, n=36). EPSP amplitudes were reduced to 66% of control by isoflurane. eEPSP rise time significantly decreased (control=4.1ms, Iso=3.3ms, P=0.038, n=33). On average, rise time decreased to 80% of control under isoflurane.

Responses of Layer 5 neurons (stimulated via both pathways) were significantly altered by isoflurane application. Pyramidal neurons in layer 5, had significantly reduced eEPSPs amplitudes under isoflurane (control=3.29mV, Iso= 1.68mV, P<0.001, n=46). Amplitudes decreased to 51% of control under isoflurane. Rise time was reduced (control=4.8ms, Iso=3.5ms, P<0.001, n=38) to 73% of control, and latency was increased (control=3.8ms, Iso=4.2ms, P=0.005, n=40) to 111% of control under isoflurane.

A linear mixed effects model was used to compare the effects of drug, pathway, and layer. There was no significant three-way interaction between isoflurane, layer and pathway, nor a significant two-way interaction between isoflurane and layer. Although, when looking at a reduced model, there was a significant effect of isoflurane on TC eEPSPs [F(1,268)=20.5, p=9.1356e-06], and on CC eEPSPs [F(1,268)=111.6, p=4.9332e-22]; as well as a significant difference in the effect of isoflurane on CC versus TC EPSPs [F(1,268)=14.2, p=2.0054e-04]. No significant correlation between other aspects of eEPSPs like latency and rise time between pathway and layer were found.

To assess the relationship between the effect of the isoflurane on input resistance in relationship to eEPSP amplitude, the regression was calculated in each layer and pathway. Interestingly, only the cortical stimulation to layer 2/3 **(figure 6A)** had a significant relationship of the effect of isoflurane on the amplitude of the eEPSP in relation to the individual neuron’s input resistance (P=0.00005). In the other layer/pathway configurations there was not a significant relationship between an individual neuron’s input resistance to its change in eEPSP amplitude by isoflurane **(figure 6B-D)**. Correlations between the neuron’s anatomical location within the depth of the cortex, with changes in eEPSP amplitude induced by isoflurane were identified. Cortical stimulation to layer 2/3 had a significant change in the correlation between cortical depth/layer to change in eEPSP amplitude with isoflurane application (P=0.01), while it was not related within the other layer/pathway configurations **(figure 6E-F)**.

**Figure 6.**
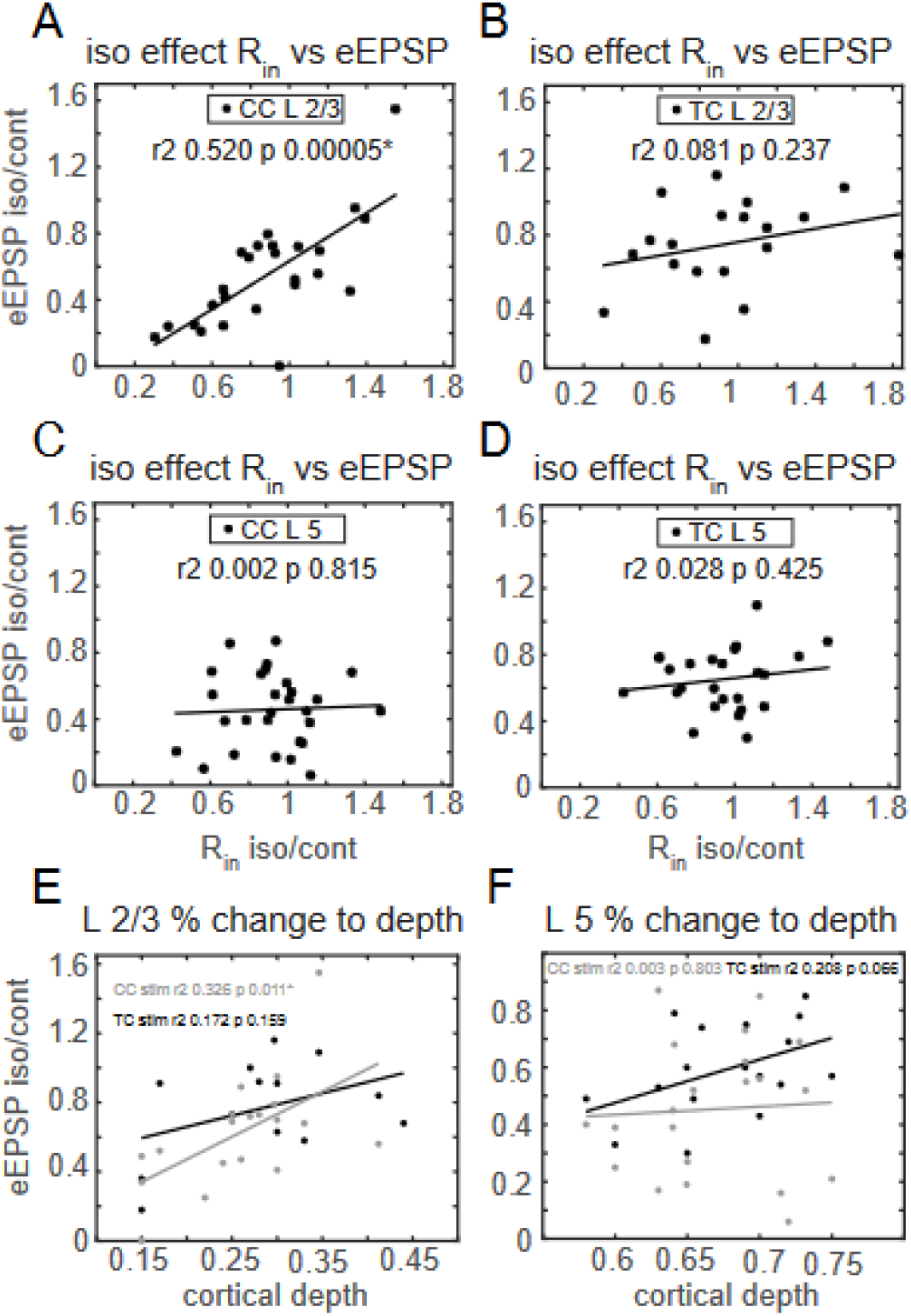
Isoflurane’s Effect on eEPSP in Relation to Input Resistance and Cortical Depth. The ratio of eEPSP amplitude recorded in isoflurane, over eEPSP amplitude in control (on the y axis), plotted against the ratio of input resistance in isoflurane, over input resistance in control (on the x axis). (A)Cortical stimulation to layer 2/3 pyramidal neurons revealed a significant relationship of eEPSP amplitude affected by isoflurane in relation to input resistance affected by isoflurane. (B)Thalamic stimulation to layer 2/3 pyramidal neurons did not reveal a correlation between eEPSP amplitude to input resistance as affected by isoflurane. (C)Cortical stimulation to layer 5 pyramidal neurons did not reveal a correlation between eEPSP amplitude to input resistance affected by isoflurane. (D)Thalamic stimulation to layer 5 pyramidal neurons did not reveal a correlation between eEPSP amplitude to input resistance affected by isoflurane. (E)Ratio of amplitude of eEPSP measured in isoflurane over control in relation to cortical depth. Layer 2/3 pyramidal neurons stimulated from the cortical pathway (gray) had a significant reduction of eEPSP in relation to cortical depth, while those stimulated by the thalamic (black) pathway did not. (F)Ratio of amplitude of eEPSP measured in isoflurane over control in relation to cortical depth. Layer 5 pyramidal neurons did have a not significant reduction of eEPSP amplitude by isoflurane that was correlated to cortical depth in either thalamic (black) stimulation, nor cortical (grey) stimulation.

Synaptic release properties were examined using paired-pulse ratio (PPR) as a demonstration of synaptic facilitation and depression. Overall, isoflurane had no significant effect on PPR. PPR was calculated as the amplitude of eEPSP peak 2/peak 1, after averaging all 11 sweeps. **Figure 7 (A-D)** shows the PPR for each layer and pathway during control, isoflurane and washout, with individual neurons illustrated in gray, and the average shown in black. Additionally, trends in the average eEPSP amplitudes of all 4 stimuli in the train, were assessed in all 3 neuronal types, with both stimulation pathways, shown in **figure 7 (E-G)**. Responses to CC stimulation in Layer 5 pyramidal neurons demonstrated depression which disappeared with isoflurane and reappeared after washout, suggesting a pre-synaptic mechanism.

**Figure 7.**
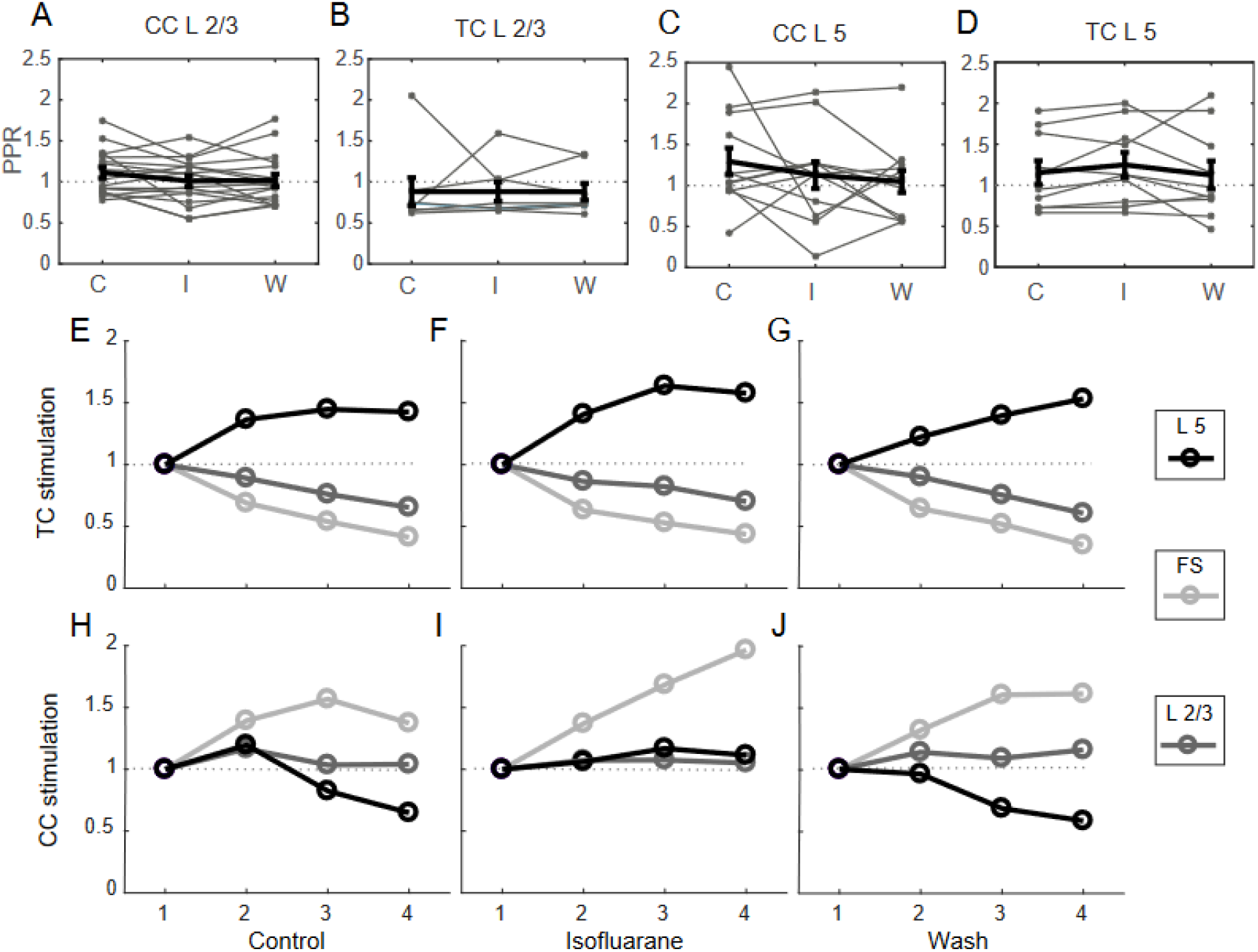
Paired Pulse Ratio & Facilitation or Depression of eEPSP Amplitude. Paired pulse ratio (PPR) was measured as the ratio of the amplitude of the second eEPSP over the amplitude of the first eEPSP from the train of eEPSPs. No significant changes occurred in PPR between control, isoflurane or wash saline in any of the pathways of stimulation or layers of pyramidal neurons recorded. Trend of synaptic facilitation and depression in each configuration based on the average amplitude is shown. (A-D) (A) PPR of L1 CC stimulation to neurons in L 2/3; (B) TC stimulation to neurons in L 2/3; (C) L1 CC stimulation to neurons in L 5; (D) TC stimulation to neurons in L 5. Black is average of all the individual traces, which are shown uniquely in gray, with no significant changes by isoflurane. (E-J) TC stimulation to layer 5 pyramidal neurons were mildly facilitating in control (E), isoflurane (F) and wash (G) saline conditions; TC onto layer 2/3 pyramidal neurons and fast spiking interneurons were mildly depressing from thalamic stimulation. (H-J) CC stimulation to fast spiking interneurons were facilitating in control (H), isoflurane (I) and wash (J) saline conditions; CC to layer 2/3 pyramidal neurons were neither facilitating or depressing, and CC to layer 5 pyramidal neurons were depressing in control and wash saline, which was relieved by isoflurane.

## Discussion

Current models of sensory awareness depict the importance of the integration of ascending and descending information streams^13-15^. One of these recently suggested models is the *information integration theory of consciousness*^16^. According to this model anesthetics act across wide areas of cortex to reduce the repertoire of network states (information) and connectivity (integration)^17,18^. Another model, derived from *predictive coding models*, suggests that the cortex is performing a comparison of incoming information to expectations mediated by top-down projections^14,19^. According to this model, loss of consciousness (LOC) would be caused by a preferential suppression of top-down information flow, preventing this comparison^20,21^. Thus, sensory information arrives intact in cortex, but top-down predictive information is absent, and therefore meaning cannot be assigned to the incoming information.

Several lines of evidence suggest that cortico-cortical connectivity is disrupted upon LOC^17,20-22^. Cortical feedback, top-down connectivity is particularly sensitive to anesthetics at hypnotic doses^9,20,21,23-28^. Furthermore, general anesthetics eliminate contextual modulation of responses in primary visual cortex that are likely mediated by descending cortico-cortical connections but leave ascending responses intact^29^, and suppress integration of local receptive field information^30^.

This work demonstrates that isoflurane anesthesia preferentially suppressed cortically evoked EPSPs over thalamically evoked EPSPs in pyramidal neurons and in fast spiking interneurons in both layer 2/3 and layer 5 of the auditory cortex. Using a linear mixed effect model, the preferential suppression of synaptic currents from top-down connectivity was greater than the bottom-up connectivity.

Other features of the eEPSPs showed common trends between the thalamic and cortical stimulation synapsing onto pyramidal neurons. Additionally, the latency to the eEPSP following cortical stimulation, but not thalamic stimulation was significantly reduced by isoflurane aesthesia. Furthermore, the rise time of the eEPSP following cortical stimulation, but not the thalamic stimulation, was reduced by isoflurane in both layers. Interestingly, none of the eEPSP features in fast spiking interneurons were significantly altered by isoflurane except the reduced eEPSP amplitude from cortical, but not thalamic, stimulation.

Intrinsic neuronal properties also showed variability between layers in response to isoflurane aesthesia. Spike threshold in both layer 2/3 and layer 5 pyramidal neurons in the auditory cortex was significantly depolarized by isoflurane, while fast-spiking interneurons did not show this change. Interestingly, layer 2/3 pyramidal neurons had a significantly shorter latency to initiation of action potential following current injection while under isoflurane aesthesia. Moreover, only layer 5 pyramidal neurons had significantly reduced input resistance when exposed to isoflurane. These changes were not reflected in fast-spiking interneurons.

In addition, when looking at paired pulse ratios following trains of stimulation, the layer 5 pyramidal neurons activated by cortical stimulation showed a distinctive response to isoflurane application. All other cell types and pathways of stimulation had consistent patterns of activation, with either depression or facilitation of the sequential eEPSPs in the train of four stimuli. The layer 5 pyramidal neurons driven cortically had an inversion of typified response to isoflurane application that was not seen in any of the other cell stimulation combinations.

Prior work has demonstrated a preferential effect of isoflurane anesthesia on the feedback, cortico-cortical pathways over the feed-forward thalamo-cortical pathways^5,6^. These results were obtained using population local field potential responses, allowing registration of a global response of the cortical columns. These results were extended to the EPSP responses of specific neurons, allowing quantification of the effects of isoflurane on the post-synaptic potentials of specific neurons. The change of EPSP amplitude paralleled the change previously reported in columnar response and may explain what underlies it.

Furthermore, our results hint to the underlying cellular mechanisms. The specific effect of isoflurane on the paired pulse ratio of the layer 5 pyramidal neurons to CC stimulation suggests a pre-synaptic effect in this type of synapse. On top of this, in was found that in layer 2/3, the EPSP amplitude suppression by isoflurane was correlated to the change in membrane resistance as well as to the distance of the neurons from the pia. This suggests an effect on the apical dendrites: effect that is related to the dendritic length (the shorter the dendrite – the stronger the effect) and is reflected in the membrane resistance (with a shorter dendrite, changes of the dendritic resistance may be reflected in the membrane resistance measured at the soma). Layer 5 neurons have longer dendrites, and changes in the apical dendrites are not reflected in the membrane resistance recorded at the soma. In this layer, all dendrites are of relatively similar length (governed mostly by the distance of the layer from the pia, rather than the location within the layer). Thus, for this group of neurons a correlation of depth was not found. The effect of isoflurane on the response to TC projections, which terminate mostly on proximal dendrites and soma, is also independent of the dendritic length.

This work concentrates on the effects of isoflurane on a specific brain region.

The results seem to support the idea of predictive coding (at least in the sensory cortex) and suggest that the relative sensitivity of the CC projections is related to its effect via apical dendrites. However, to prove that the dendritic effect is indeed the reason for this selective effect, one must perform direct recording from apical dendrites. Furthermore, to convince that this is indeed the mechanism underlying anesthetic induced disconnectedness, examination of whether this effect can be generalized to other brain regions must be made, along with other anesthetic drugs that affect different molecular targets.

## Conclusion

Isoflurane differentially suppresses top-down synaptic transmission preferentially over bottom-up afferent sensory communication. Unique eEPSP characteristics are modulated between layer and pathway by isoflurane aesthesia. Intrinsic neuronal properties were distinctively altered by isoflurane depending on cell body layer within the auditory cortex. Furthermore, response features of fast spiking interneurons, although not recorded in large numbers were not modulated by isoflurane aesthesia like pyramidal neurons, other than the cortically evoked EPSP amplitude reduction. Neuronal modulation by isoflurane was exclusive to cell type, layer and origin of the stimulation.

## Acknowledgements

Supported by the National Institutes of Health (Bethesda, MD) DC006013 (to M. I. Banks), Mentored research award grant from the International Anesthesia Research Society (to A. Raz), a grant from the United States - Israel Binational Science Foundation (BSF), Jerusalem, Israel (to A. Raz) and the Department of Anesthesiology, School of Medicine and Public Health, University of Wisconsin, Madison, WI. The authors are grateful for the assistance of Dr. Bryan Krause with the statistical analysis. The authors also thank Sean Grady for technical support on this project.

---

**Table 1.** Intrinsic Properties of Layer 2/3 pyramidal neurons.

| Variable | Control | Isoflurane | Wash | n | P value |
| --- | --- | --- | --- | --- | --- |
| Resting Membrane | -76.19 mV | -74.88 mV | -74.31 mV | 28 | 0.064 |
| Spike Threshold | -36.85 mV | -33.62 mV | -35.78 mV | 26 | **0.003** |
| Input Resistance | 115.96 MΩ | 117.55 MΩ | 122.23 MΩ | 28 | 0.558 |
| Spike Latency | 90.2 msec | 55.0 msec | 66.8 msec | 25 | **0.004** |
| Time Constant | 9.3 msec | 10.3 msec | 10.7 msec | 26 | 0.174 |
| TC stim | 4.40 mV | 3.37 mV | 5.04 mV | 15 | **0.015** |

**Table 2.** Intrinsic Properties of Layer 5 pyramidal neurons.

| Variable | Control | Isoflurane | Wash | n | P value |
| --- | --- | --- | --- | --- | --- |
| Resting Membrane | -68.85 mV | -68.38 mV | -68.27 mV | 33 | 0.693 |
| Spike Threshold | -38.57 mV | -34.80 mV | -36.98 mV | 33 | **< 0.001** |
| Input Resistance | 142.82 MΩ | 123.52 MΩ | 140.96 MΩ | 33 | **0.013** |
| Spike Latency | 89.8 msec | 86.2 msec | 77.0 msec | 33 | 0.411 |
| Time Constant | 15.0 msec | 15.3 msec | 15.9 msec | 33 | 0.567 |

**Table 3.** Intrinsic Properties of Fast Spiking interneurons.

| Variable | Control | Isoflurane | Wash | n | P value |
| --- | --- | --- | --- | --- | --- |
| Resting Membrane | -70.89 mV | -72.84 mV | -72.51 mV | 8 | 0.276 |
| Spike Threshold | -37.07 mV | -37.21 mV | -38.71 mV | 8 | 0.525 |
| Input Resistance | 119.45 MΩ | 113.91 MΩ | 124.82 MΩ | 8 | 0.198 |
| Spike Latency | 93.0 msec | 60.2 msec | 76.7 msec | 8 | 0.339 |
| Time Constant | 5.4 msec | 6.0 msec | 6.2 msec | 8 | 0.533 |

**Table 4.** eEPSP characteristics in Layer 2/3 pyramidal neurons.

| Variable | Control | Isoflurane | Wash | n | P value |
| --- | --- | --- | --- | --- | --- |
| TC stim | 4.40 mV | 3.37 mV | 5.04 mV | 15 | **0.015** |
| CC stim | 5.32 mV | 3.21 mV | 5.44 mV | 21 | **< 0.001** |
| PPR TC | 0.88 | 0.88 | 0.88 | 8 | 0.993 |
| PPR CC | 1.11 | 1.01 | 1.02 | 17 | 0.135 |
| Rise time TC | 2.8 msec | 2.5 msec | 3.0 msec | 13 | 0.573 |
| Rise time CC | 5.0 msec | 3.8 msec | 4.3 msec | 20 | **0.021** |
| Latency TC | 2.4 msec | 2.5 msec | 2.5 msec | 15 | 0.426 |
| Latency CC | 3.7 msec | 3.8 msec | 3.8 msec | 20 | 0.738 |

**Table 5.** eEPSP characteristics in Layer 5 pyramidal neurons.

| Variable | Control | Isoflurane | Wash | n | P value |
| --- | --- | --- | --- | --- | --- |
| TC stim | 1.82 mV | 1.16 mV | 1.80 mV | 20 | **< 0.001** |
| CC stim | 4.42 mV | 2.08 mV | 4.26 mV | 26 | **< 0.001** |
| PPR TC | 1.15 | 1.25 | 1.12 | 10 | 0.365 |
| PPR CC | 1.29 | 1.13 | 1.05 | 12 | 0.301 |
| Rise time TC | 3.5 msec | 2.8 msec | 3.6 msec | 15 | 0.050 |
| Rise time CC | 5.7 msec | 4.0 msec | 5.4 msec | 23 | **< 0.001** |
| Latency TC | 3.3 msec | 3.6 msec | 3.4 msec | 17 | 0.132 |
| Latency CC | 4.2 msec | 4.7 msec | 4.3 msec | 23 | **0.029** |

**Table 6.** eEPSP characteristics in Fast Spiking interneurons.

| Variable | Control | Isoflurane | Wash | n | P value |
| --- | --- | --- | --- | --- | --- |
| TC stim | 5.65 mV | 4.99 mV | 8.05 mV | 6 | 0.172 |
| CC stim | 6.88 mV | 3.81 mV | 6.94 mV | 8 | **< 0.001** |
| PPR TC | 1.18 | 1.25 | 1.02 | 6 | 0.333 |
| PPR CC | 1.25 | 1.55 | 1.94 | 8 | 0.539 |
| Rise time TC | 1.5 msec | 1.3 msec | 1.6 msec | 6 | 0.342 |
| Rise time CC | 2.6 msec | 2.6 msec | 3.4 msec | 7 | 0.053 |
| Latency TC | 2.6 msec | 2.6 msec | 2.7 msec | 6 | 0.299 |
| Latency CC | 3.9 msec | 4.5 msec | 3.9 msec | 8 | 0.098 |

**Table 7.**
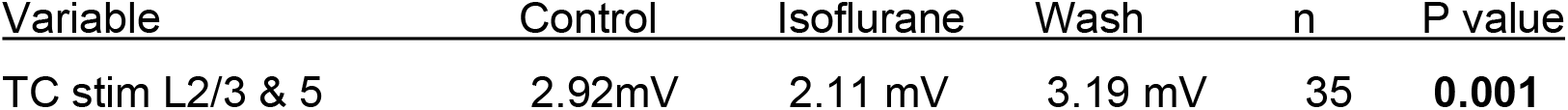
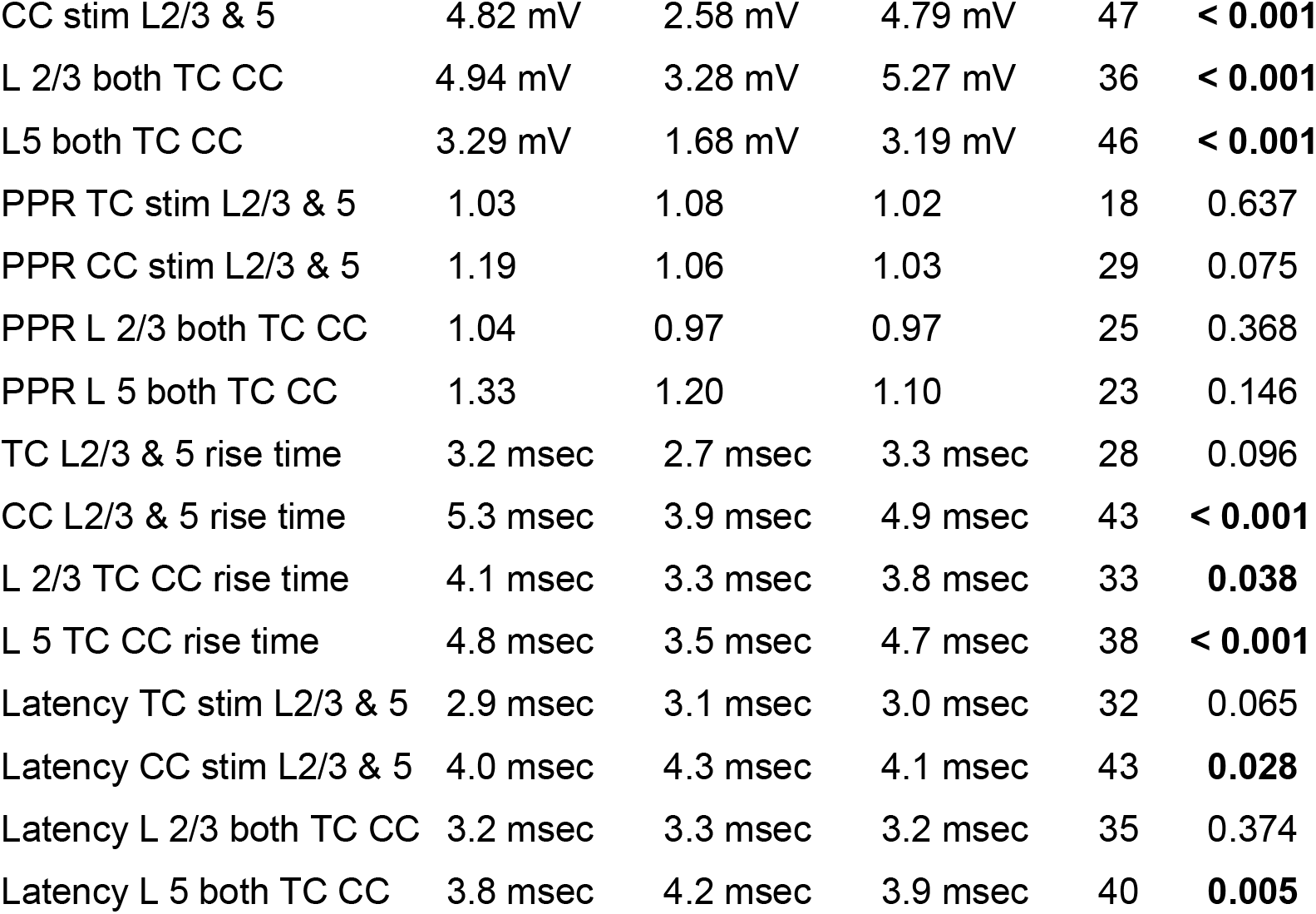
Pathway vs. layer eEPSP comparisons in pyramidal neurons.

